# Quaternary Ammonium Salt-Modified Isabgol (Psyllium) Scaffold as an Antibacterial Dressing for Improved Wound Healing

**DOI:** 10.1101/2022.04.24.489269

**Authors:** T K Vasudha, Anand Kumar Patel, Vignesh Muthuvijayan

## Abstract

Chronic wounds require suitable treatment and management strategies for proper healing. Among other causes, infection delays the healing of wounds and increases the risk of wound-related complications. Healing of chronic wounds requires an ingenious biomaterial that is biocompatible and anti-infective to achieve effective wound management. In this study, a wound dressing with inherent antibacterial and biocompatible properties was developed to assist the healing process. Natural polysaccharide Isabgol was chemically modified with Epoxy propyl trimethyl ammonium chloride to render antibacterial activity to the material. This is the first report of such chemical modification of this polymer for biomedical applications. The modified material was freeze-dried to obtain scaffolds. ^13^C NMR and FTIR analysis confirmed the modification of the Isabgol polymer chains with EPTMAC. The scaffold exhibits an organized porous structure that allows the exchange of gases and nutrients through the matrix, as confirmed by SEM analysis. The material possesses excellent swelling properties up to 17 times its initial weight that allows it to absorb wound exudates and maintain a moist environment at the wound site. The scaffold is biodegradable, and thermally and mechanically stable. The material is anti-infective and can prevent infections at the wound site, which is one of the major causes of delayed wound healing. The developed scaffolds have been proven to be biocompatible and suitable for use in blood contact applications. Finally, since Isabgol is a low-cost raw material, the quaternary ammonium-modified Isabgol scaffold can be an affordable wound dressing material.

## 1. INTRODUCTION

Wound healing is an orderly progression of biological and molecular events that occurs in roughly four stages, namely hemostasis, inflammation, proliferation and remodelling stages. In the hemostasis phase, platelets arrive at the wound site and clot formation occurs to cover the wound and ward-off bacteria. In the inflammation stage, wound debridement action is performed by macrophages and neutrophils that are recruited to the wound site. These inflammatory cells remove cell debris and bacteria at the wound site. The proliferation stage is characterized by the formation of ECM proteins, angiogenesis, wound contraction, and keratinocyte migration. In the remodelling phase, collagen fibres deposited in the proliferative phase are aligned in an orderly fashion to increase the tensile strength of the newly formed tissue[1]. Infection is one of the leading causes of impaired wound healing. Infection results in prolonged inflammation where the accumulation of inflammatory cells causes the production of inflammatory cytokines that secrete matrix metalloproteases (MMPs) that destroy the wound healing process. Additionally, production of factors like vascular endothelial growth factor (VEGF) and platelet-derived growth factor (PDGF) is also affected. Under these conditions, the transition from the inflammatory to proliferative stage does not occur hence preventing the formation of new healthy tissue. It is, therefore, crucial to prevent infection at the site of the wound [1-3].

To prevent wound infection and encourage wound healing, an appropriate wound dressing material can be used to assist various aspects of the wound healing process. An ideal wound dressing material is biocompatible, non-toxic, provides a moist wound environment, promotes angiogenesis, allows gas and nutrient exchange, and promotes cell adhesion, proliferation, migration, and cell differentiation[4]. Various standalone and combinations of natural and synthetic biomaterials such as collagen[5, 6], chitosan[7, 8], alginate[9, 10], silk fibroin [11, 12], poly (ethylene glycol)[13, 14], poly (vinyl alcohol)[15, 16], poly (lactic-co-glycolic acid)[17, 18] etc. have been explored for use as wound dressing materials. Although several commercially available materials have been developed to assist the wound healing process, they are prohibitively expensive and are not affordable for most patients[19]. In the present study, we have developed an ingenious and inexpensive wound dressing material that is biocompatible and anti-infective to achieve effective wound management.

We have used Isabgol (psyllium), a natural plant polysaccharide containing xylose and arabinose monomers in our wound dressing. Isabgol contains an alkali-soluble and an alkali-insoluble fraction. The alkali-soluble fraction (∼85%) is the arabinoxylan mucilage polysaccharide that gels over a range of concentrations and is of interest for the wound dressing application. Isabgol is regarded safe for ingestion and is largely used to treat bowel disorders[20]. Isabgol is highly hydrophilic and is capable of absorbing wound exudates. It provides a moist environment that helps to accelerate wound healing. Isabgol has been proven to be biocompatible and suitable for biomedical applications[21, 22]. It is an inexpensive polysaccharide source, therefore the final wound dressing material that we have obtained will be available at an affordable price in the market.

Quaternary ammonium compounds (QAC) are cationic molecules possessing potent antimicrobial properties. They are of the general structure N^+^R1R2R3R4X^-^ where X is an anion and R is a hydrogen atom or an alkyl group. QACs have been useful in disinfectants, pharmaceutical, hygiene and healthcare products for their antimicrobial action, which has been mainly attributed to the electrostatic interaction between the positively charged quaternary nitrogen of QAC and negatively charged phospholipids in bacterial membranes. 2,3-Epoxypropyltrimethylammonium chloride (EPTMAC) is a QAC that has been used in various applications including antibacterial textiles[23, 24], food packaging[25, 26], and biological applications[27, 28]. We have previously been successful in demonstrating the use of Isabgol in biomedical applications, especially for wound healing[21, 22]. In this study, we have chemically modified Isabgol polymer with EPTMAC to impart antimicrobial properties to the wound dressing material to prevent infection at the wound site and accelerate the wound healing process. We have studied the physicochemical properties of the material and performed biological evaluation of the biomaterial to assess its suitability for wound healing applications.

## 2. MATERIALS AND METHODS

### 2.1 Synthesis of Isabgol and EPTMAC-Modified Isabgol Wound Dressings

Isabgol was purchased at a local store in Chennai, India. EPTMAC was purchased from Sigma-Aldrich. 2 (w/v) % Isabgol solution was prepared in 2 (w/v) % Sodium hydroxide. The solution was kept at 55-60 °C for 5 hours under constant stirring condition. After Isabgol dissolved completely, the solution was filtered to remove the alkali-insoluble fraction. 2 (v/v) % EPTMAC was added to the filtered Isabgol solution. The reaction was allowed to occur at 55 ° C for 2.5 hours at constant stirring condition. Both the solutions, pure Isabgol (ISAB) and modified Isabgol (MISAB) were dialysed against reagent grade water under stirring condition for 48 hours. The hydrogel obtained was then frozen at -80 °C for 12 hours and lyophilized for 48 hours to obtain porous scaffolds.

### 2.2 Chemical Characterization

^13^C NMR spectra of ISAB, MISAB, and EPTMAC were recorded using Bruker AVANCE HD 500MHz FT-NMR Spectrometer. Additionally, to confirm that EPTMAC has been incorporated into ISAB by chemical crosslinking and the final product is not simply a physical mixture of Isabgol and EPTMAC, a physical blend of EPTMAC with Isabgol was prepared in the absence of NaOH and was subjected to ^13^C NMR after 48 hours of dialysis against distilled water. The functional groups on ISAB and MISAB were studied by Fourier Transform Infrared Spectroscopy (FTIR) in the attenuated total reflectance (ATR) mode. The degree of substitution of EPTMAC on Isabgol was evaluated by studying the elemental composition of ISAB and MISAB using a CHNS/O PerkinElmer 2400 series analyser.

### 2.3 Physical Characterization

#### 2.3.1 Scanning electron microscopy

The surface morphology of the synthesized scaffolds was studied using scanning electron microscopy (SEM). The ISAB and MISAB hydrogels were freeze-dried and sputtered with gold followed by SEM imaging. The porosity of the materials was calculated using ImageJ software.

#### 2.3.2 Swelling Kinetics

Swelling kinetics study was performed to determine the degree of swelling of the scaffolds at equilibrium. Briefly, ISAB and MISAB scaffolds of approximately equal weight and volume were immersed in PBS. Immersed samples were taken out and their weight was measured at regular time intervals until the samples began to disintegrate.

The swelling ratio was calculated using the following formula:

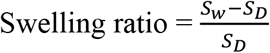

where S_D_ is the sample dry weight and SW is the sample wet weight.

#### 2.3.3 Thermogravimetric analysis

The thermal stability of ISAB and MISAB was evaluated using the thermogravimetric analyser (Agilent TGA-DTA instrument) in nitrogen atmosphere. The study was carried out from 25°C to 800 °C at a ramp rate of 10 °C/min.

#### 2.3.5 Mechanical behaviour

The compressive strength and modulus of the scaffolds were determined by uniaxial compression testing using BISS UTM-5 kN at a compression rate of 0.5 mm/min. Scaffolds of 20 × 20 × 7 mm dimensions were used and the measurements were conducted at room temperature. The compression modulus was calculated using Hooke’s law.

### 2.4 Biological Characterization

#### 2.4.1 Antibacterial activity

Antibacterial activity of the ISAB and MISAB scaffolds was evaluated using both gram-negative (*E. coli*) and gram-positive (*S. aureus*) bacteria. Bacterial cultures were grown till their log phase was reached and the cultures were incubated with the scaffolds (20 mg ml^-1^) for 8 hours. Bacterial culture with no scaffold incubated in it was treated as negative control. After incubation, the antibacterial activity was measured by counting the number of colonies formed per mL (CFU/mL).

#### 2.4.2 Hemocompatibility assay

To demonstrate the compatibility of ISAB and MISAB for blood contact applications such as wound management, the material leachates were incubated in a suspension of RBCs for 8 hours. Triton X-100 was added as the positive control and PBS as the negative control. After the incubation, the RBC suspension was centrifuged. The absorbance of the supernatant at 540 nm was measured. Hemolysis percentage was interpreted as follows: <2%: non-hemolytic, 2– 5%: slightly hemolytic and >5% hemolytic samples [29]. The RBCs in the pellet were fixed with 1% glutaraldehyde and their morphology was assessed by SEM imaging. All the protocols pertaining to the use of blood samples were approved by the Institute Ethics Committee, IIT Madras (IEC/2020-03/VM/17).

Percentage hemolysis was calculated as follows:

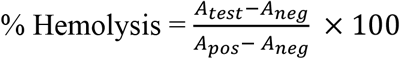

Where *A*_*test*_ is the absorbance of the RBC suspension incubated with scaffold leachate.

*A*_*pos*_ is the absorbance of the RBC suspension incubated with Triton X-100 (positive control)

*A*_*neg*_ is the absorbance of the RBC suspension incubated with PBS (negative control)

#### 2.4.3 Cell culture studies

3T3-L1 mouse fibroblast cells were used for the cell culture studies. The cells were maintained in DMEM culture media with 10% fetal bovine serum and 1% gentamycin-penicillin-streptomycin in an incubator at 37 °C and 5% Carbon dioxide.

##### 2.4.3.1 Cell Viability Assay

Leachates of ISAB and MISAB were prepared by extracting the material in DMEM culture medium for 24 hours. To test the cell viability of 3T3-L1 mouse fibroblasts in the presence of the leachates, a 3-(4,5-dimethylthiazol-2-yl)-2,5-diphenyl tetrazolium bromide (MTT) assay was conducted. Briefly, eighty percent confluent cells were trypsinized and seeded on to a 96-well plate (1 × 10^4^ cells/well) and incubated for 24 hours. Following incubation, the cells were treated with ISAB and MISAB leachate at 20 mg ml^-1^ concentration for 24, 48 and 72 hours. Tissue culture plate with cells without treatment with any leachate was considered negative control. After the incubation period, the contents of the well were removed, 100 µl MTT was added to the wells and the plate was incubated in the dark at 37°C and 5% carbon dioxide for 1 hour. The appearance of formazan crystals was observed after 1 hour, post which the MTT reagent was removed and Dimethyl sulfoxide (DMSO) was added to dissolve the crystals. The absorbance of the wells was measured using a microplate reader at 570 nm with 630 nm as reference wavelength. The experiment was performed in triplicate and repeated thrice. The percent cell viability was calculated as:

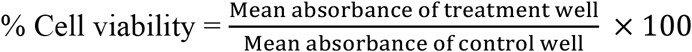

##### 2.4.3.2 Scratch Wound Assay

3T3-L1 mouse fibroblast cells were cultured in a 6-well plate. When confluence was reached, cell monolayers were gently scratched with a sterile micropipette tip and extensively rinsed with medium to remove all cellular debris. The cells were treated with ISAB and MISAB leachate in the treatment wells. The proliferation of cells into the wound area after 6, 12 and 24 hours was observed under light microscope and was photographed for examining the wound area.

##### 2.4.3.3 Live Dead Assay

3T3-L1 mouse fibroblast cells were seeded in a 12 well plate and incubated for 24 hours. The cells were then treated with ISAB and MISAB leachate for 24 hours. Following incubation, the contents of the wells were removed, washed thoroughly with sterile PBS and stained with SYTO 9/ Propidium iodide dye mixture for 15 minutes. The wells were then washed with PBS and the cells were fixed with 3.7% formaldehyde at 4°C for 10 minutes and room temperature for 10 minutes and washed again with PBS. 1 ml of PBS was added to each well. The cells were visualized under an inverted fluorescence microscope (Olympus IX83).

### 2.5 Statistical Analysis

All experiments were performed in triplicates and were repeated thrice. The values have been expressed as mean ± standard deviation. Statistical significance of the data was determined by one way ANOVA or student t-test. GraphPad Prism 7 was used to perform statistical analyses.

## 3. RESULTS AND DISCUSSION

### 3.1 Synthesis and Characterization of polymers

Isabgol hydrogels were synthesized by dissolving the alkali-soluble gelling fraction of psyllium husk in NaOH, followed by filtration and dialysis to obtain Isabgol hydrogels. The EPTMAC-modified Isabgol hydrogels (MISAB) were obtained by addition of EPTMAC to Isabgol dissolved in NaOH. The reaction of Isabgol with EPTMAC results in nucleophilic substitution and opening of the epoxide ring on EPTMAC. The hydroxyl groups on Isabgol act as the nucleophilic centres and react with the epoxide ring. Figure 1 represents the synthesis scheme for the modification of Isabgol polymer chains with EPTMAC.

**Figure 1:**
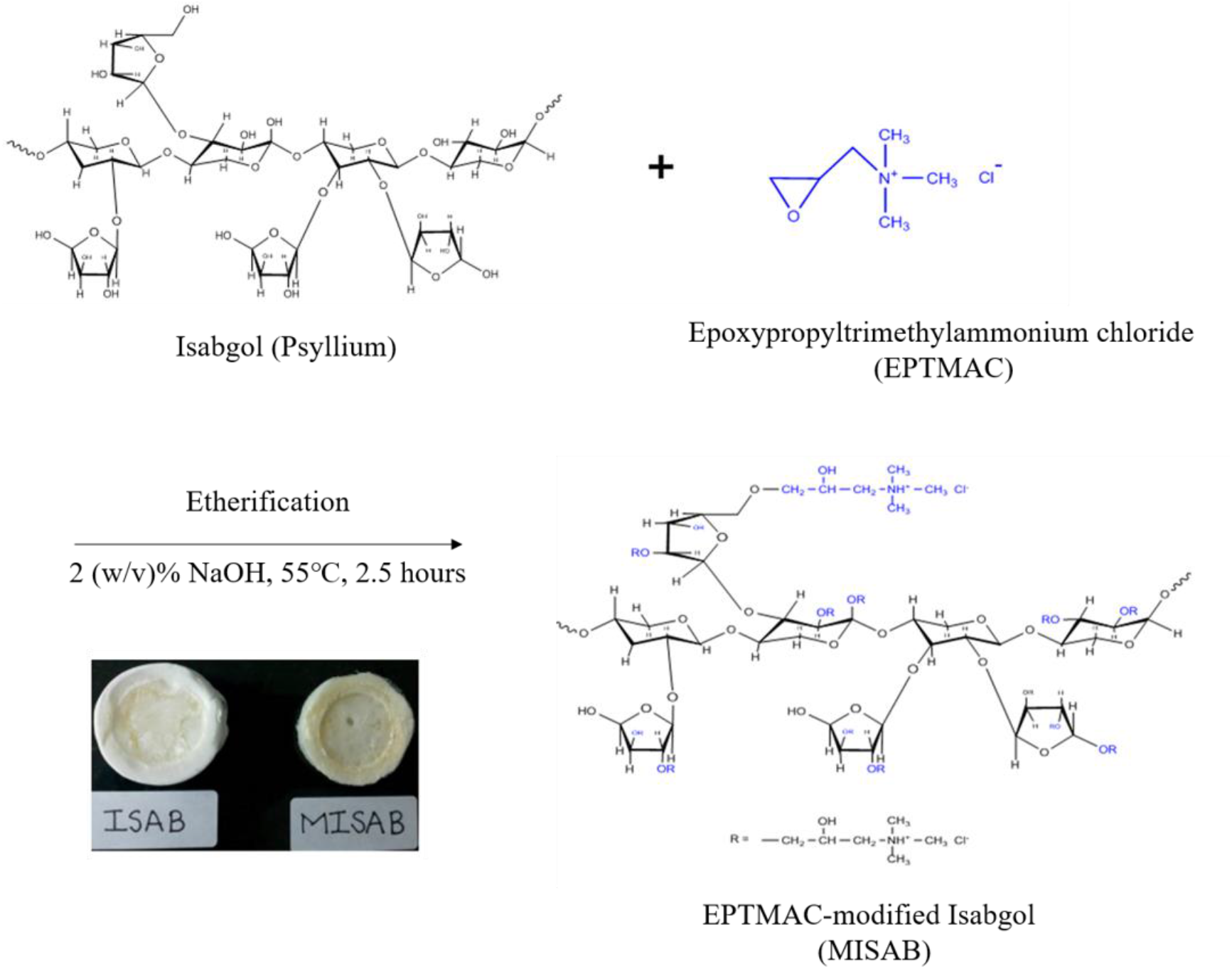
Synthesis scheme for the functionalization of Isabgol with EPTMAC

The functionalization of Isabgol polymer chains with EPTMAC was confirmed by ^13^C NMR spectroscopy and FTIR spectroscopy. 13C NMR spectrum of EPTMAC-modified scaffolds showed a characteristic peak at 55 ppm which is absent in the ^13^C NMR spectrum of pure Isabgol scaffolds (Figure 2a) [36]. The ^13^C NMR spectrum of modified Isabgol presents the same peaks as unmodified Isabgol, but they are an addition to other relative peaks to the specific carbons of the quaternary ammonium graft.

**Figure 2:**
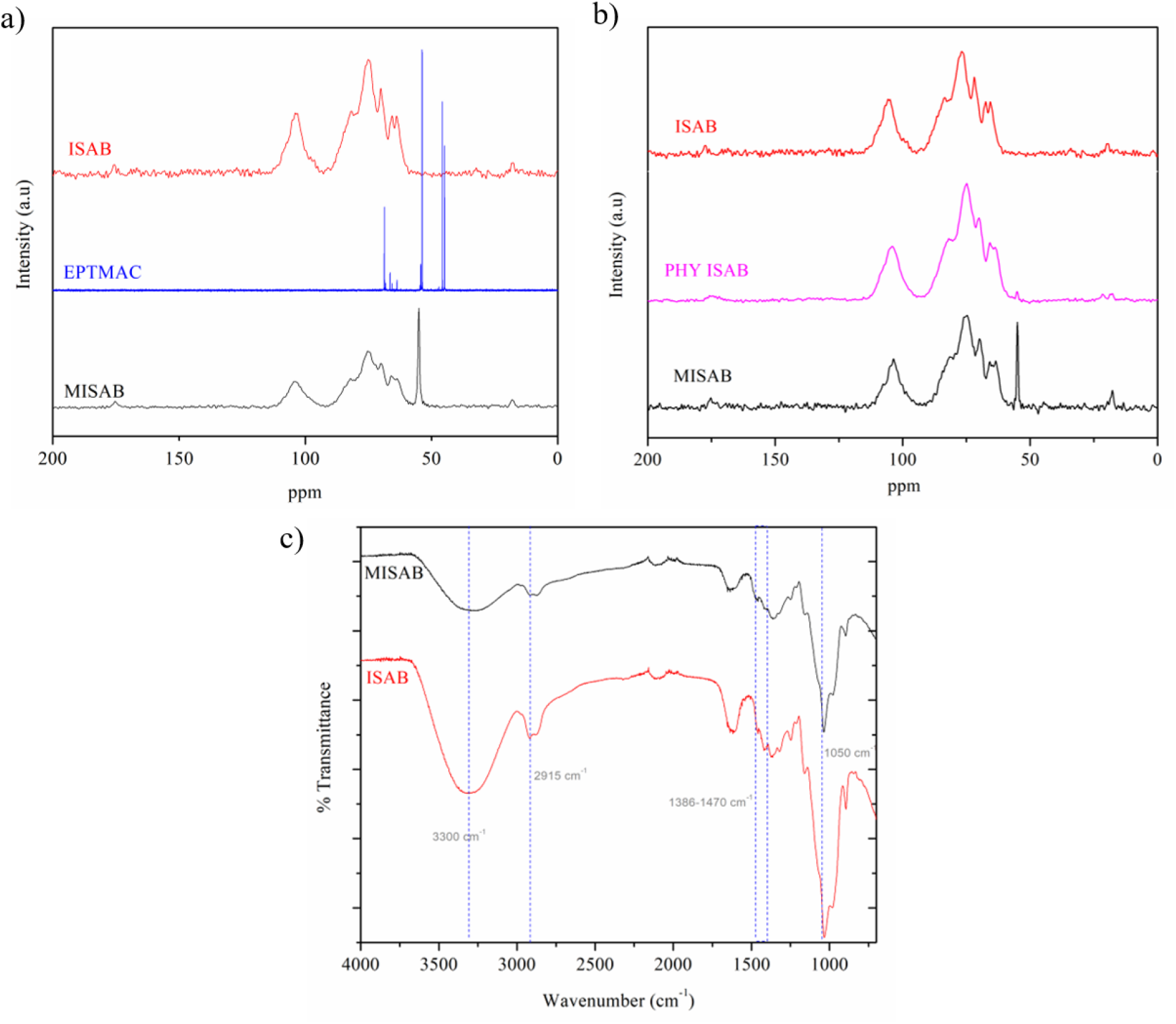
a) ^13^C NMR spectra showing EPTMAC crosslinking to Isabgol polymer chains b) ^13^C NMR spectra confirming chemical crosslinking of EPTMAC to Isabgol c) FTIR spectra of ISAB and MISAB

To confirm that the characteristic EPTMAC peak is due to chemical crosslinking of the quaternary ammonium salt and not due to a physical blend with the Isabgol, a physical blend of EPTMAC with Isabgol prepared in the absence of NaOH was subjected to NMR after 48 hours of dialysis against distilled water. The characteristic peaks at 55 ppm indicating the presence of quaternary ammonium groups were absent, which confirms that the presence of characteristic peaks for quaternary ammonium groups is due to chemical crosslinking only (Figure 2b).

The FTIR analysis was performed in the ATR (attenuated total reflectance) mode. The MISAB structure contains a unique chemical link (C–H in the methyl group) that is not present in ISAB. Figure 6 shows the FTIR spectra of ISAB and MISAB.The sharp band at 3300 cm^-1^ belongs to the (O–H) groups. Hydroxyl related peak weakened strongly for MISAB because of ring-opening reaction with epoxy groups of EPTMAC. The peaks obtained at 1386–1470 and about 2915 cm^-1^ are, respectively, due to C–H bending and stretching of methyl groups. The peak at 1050 cm^-1^ can be ascribed to the C-O-C stretch of ether. These observations corroborate the results obtained through NMR spectroscopy (Figure 2c).

The nitrogen content of the modified Isabgol was measured via elemental analysis to determine the degree of substitution (Table 1). Mass percentages of hydrogen, carbon, and nitrogen were measured. The rest of the mass was assumed to be oxygen. The degree of substitution was calculated using the given formula:

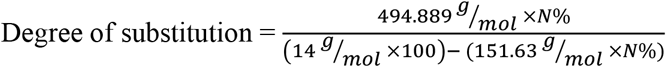

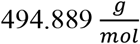 = Molecular mass of Isabgol repeat unit

**Table 1:** Elemental analysis of ISAB and MISAB polymers using CHNSO analyser

|  | %C | %H | %N | C mol% | N mol% | DS |
| --- | --- | --- | --- | --- | --- | --- |
| **ISAB** | 40.16 | 5.34 | 0.31 | 27.69 | 0.183 | - |
| **MISAB** | 23.04 | 5.25 | 2.31 | 16.45 | 1.414 | **0.59** |

N% = Nitrogen content of modified Isabgol

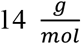= Molecular mass of nitrogen

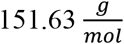= Molecular mass of EPTMAC

Higher degree of substitution of EPTMAC on Isabgol was possible. However, beyond 59% substitution of the -OH groups, hydrogen bonding between polymer chains of Isabgol was insufficient to form hydrogels. At 59% substitution, we were able to achieve a balance of developing hydrogels and obtaining desired antibacterial activity.

### 3.2 Physical characterization of ISAB and MISAB

#### 3.2.1 Scanning electron microscopy of scaffolds

The surface morphology of the synthesized scaffolds was studied using SEM imaging. The SEM images of the scaffolds reveal the porous nature of the scaffold surfaces (Figure 3a). A significant change in the surface morphology is observed after modification with EPTMAC. The microporous nature of the scaffolds can elicit the attachment of fibroblast cells, and facilitate its proliferation. The interconnected porous structure mimics the extracellular matrix rendering the scaffolds suitable for biomedical applications[30]. The porosity of the scaffolds was determined by analysing SEM micrographs of the scaffold using ImageJ software. The percentage porosity was found to be 43.7±1.4% and 51.9±2% for ISAB and MISAB scaffolds, respectively (Figure 3b). The increase in porosity in MISAB is due to the reduced number of hydroxyl groups available for crosslinking in the Isabgol polymer after modification of the hydroxyl groups with EPTMAC. Reduction in polymer crosslinking has resulted in increased porosity of the scaffold.

**Figure 3:**
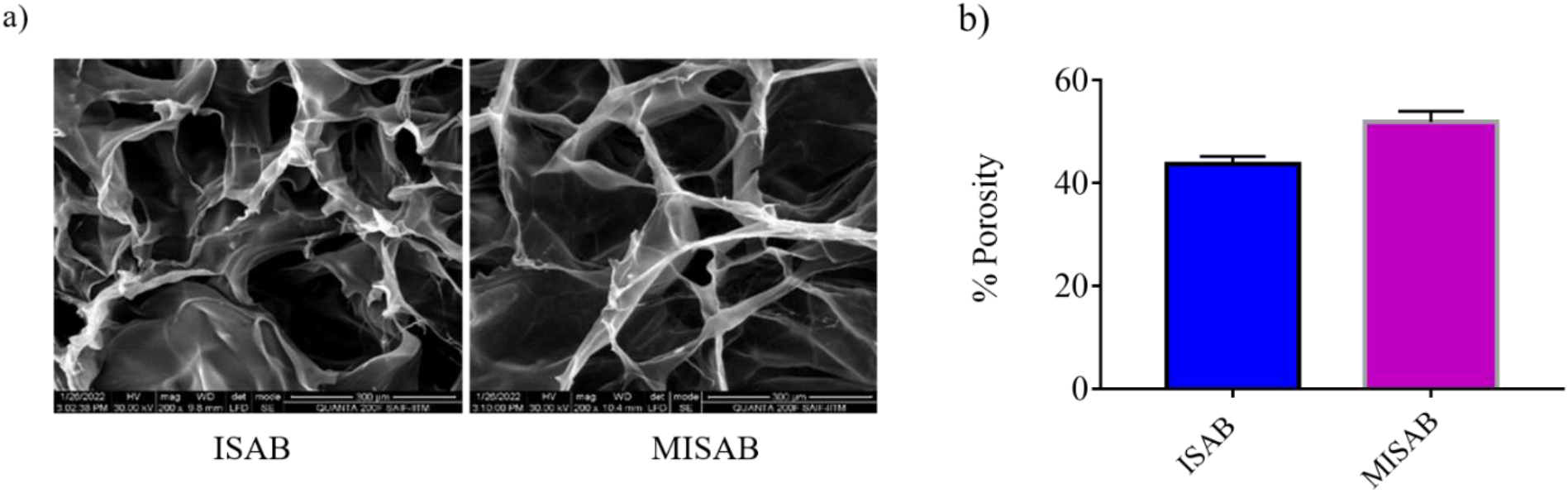
a) SEM images of ISAB and MISAB scaffolds revealing the porous nature of the scaffolds b) Percentage porosity of scaffolds estimated using ImageJ

#### 3.2.2 Swelling and *in vitro* degradation study

The swelling ability of the prepared scaffolds was tested using phosphate buffered saline (PBS) absorption assay. Both ISAB and MISAB displayed excellent swelling ability, which allows the material to absorb wound exudates ensuring a moist wound environment and prevention of bacterial contamination. The MISAB scaffolds, however, showed reduced swelling compared to the ISAB scaffolds. This may be attributed to the reduced number of — OH groups after EPTMAC modification. ISAB showed a swelling ratio of 23.3 ± 0.3, while MISAB displayed a swelling ratio of 17.4 ± 0.4 after 12 hours of incubation in PBS (Figure 4a). To study the biodegradation of the scaffolds, they were immersed in 10000 U/mg lysozyme. Lysozyme hydrolyzes the glycosidic bonds of the Isabgol polymer leading to the degradation of the scaffolds. Figure 4c reveals the loss of organized porous structure of the scaffolds after 24h of incubation in lysozyme at 37°C.

**Figure 4:**
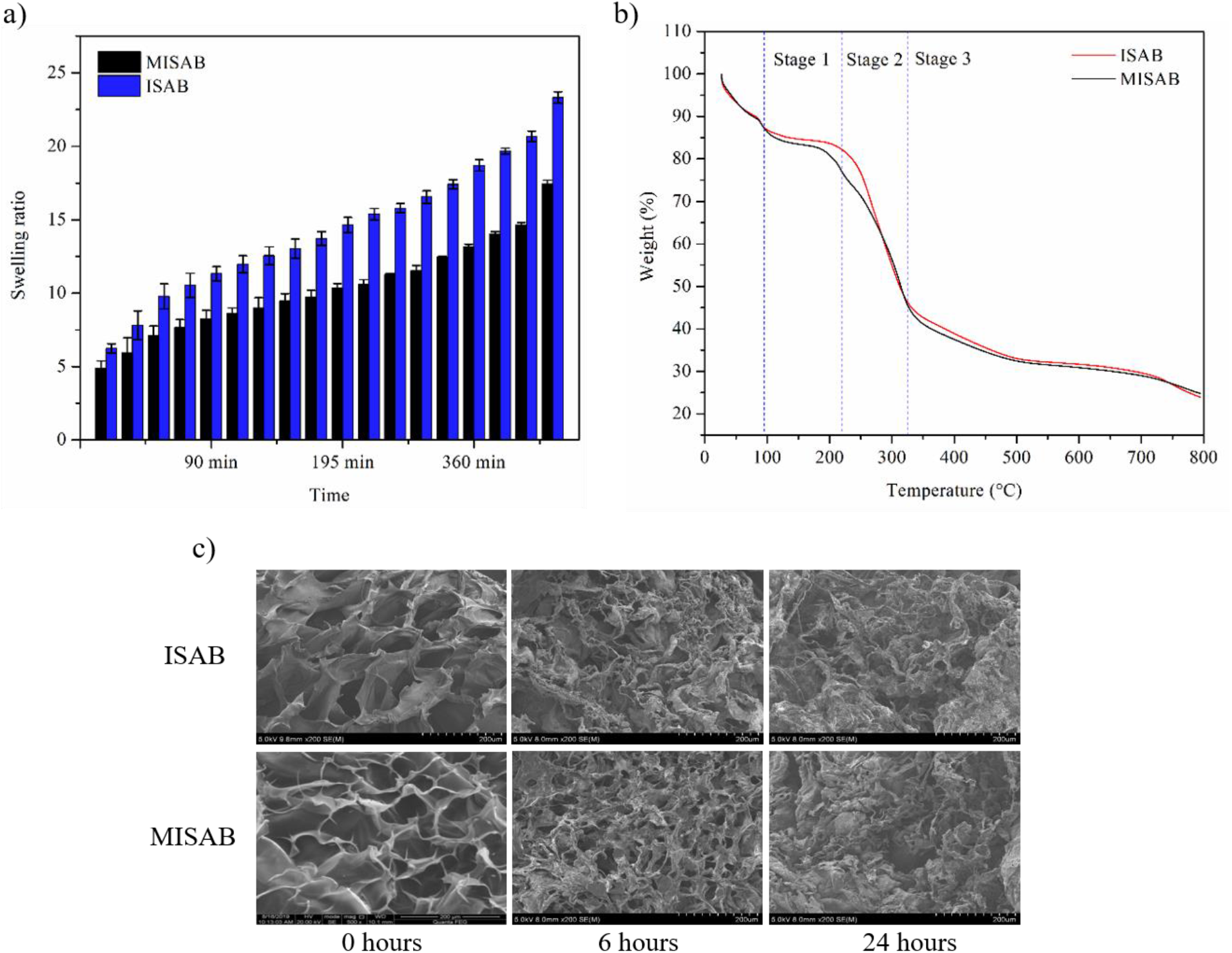
a) Swelling kinetics-PBS absorption assay b) TGA curves of ISAB and MISAB c) SEM micrographs showing the loss of organized porous structure of ISAB and MISAB scaffolds in the presence of 10000 U/mg lysozyme

#### 3.2.3 Thermal stability of scaffolds

The results of TGA of ISAB and MISAB revealed 3 stages of decomposition. The modification of Isabgol with EPTMAC did not change the thermal properties of Isabgol. The weight loss around 100 °C is due to the loss of moisture from the scaffolds. The first stage of polymer decomposition occurs in the temperature range from 100 to 220°C and corresponds to the breakdown of polymer chains resulting in 5% and 11% weight loss in ISAB and MISAB respectively. In the subsequent stages, further decomposition of residual oligo- and mono-saccharides occurs and results in the evolution of CO2 and water vapours. In stage 2 (220-324°C), weight loss of 35% (ISAB) and 30% (MISAB) was observed followed by a weight loss of 21% in both ISAB and MISAB scaffolds in stage 3 (>324°C) (Figure 4b).

#### 3.2.5 Mechanical behaviour

Compression stress-strain testing was performed on the scaffolds. The scaffolds displayed good mechanical performance. The compressive strength and the Young’s modulus of ISAB and MISAB was calculated from the stress-strain curves. The Young’s modulus was found to be 0.254±0.002 MPa and 0.258±0.005 MPa for ISAB and MISAB respectively. The compressive strength was calculated to be 1.02±0.09 MPa for ISAB and 0.907843553 ± 0.10 MPa for MISAB (Figure 5). The compressive strength of ISAB was slightly higher than that of MISAB but was not significantly different.

**Figure 5:**
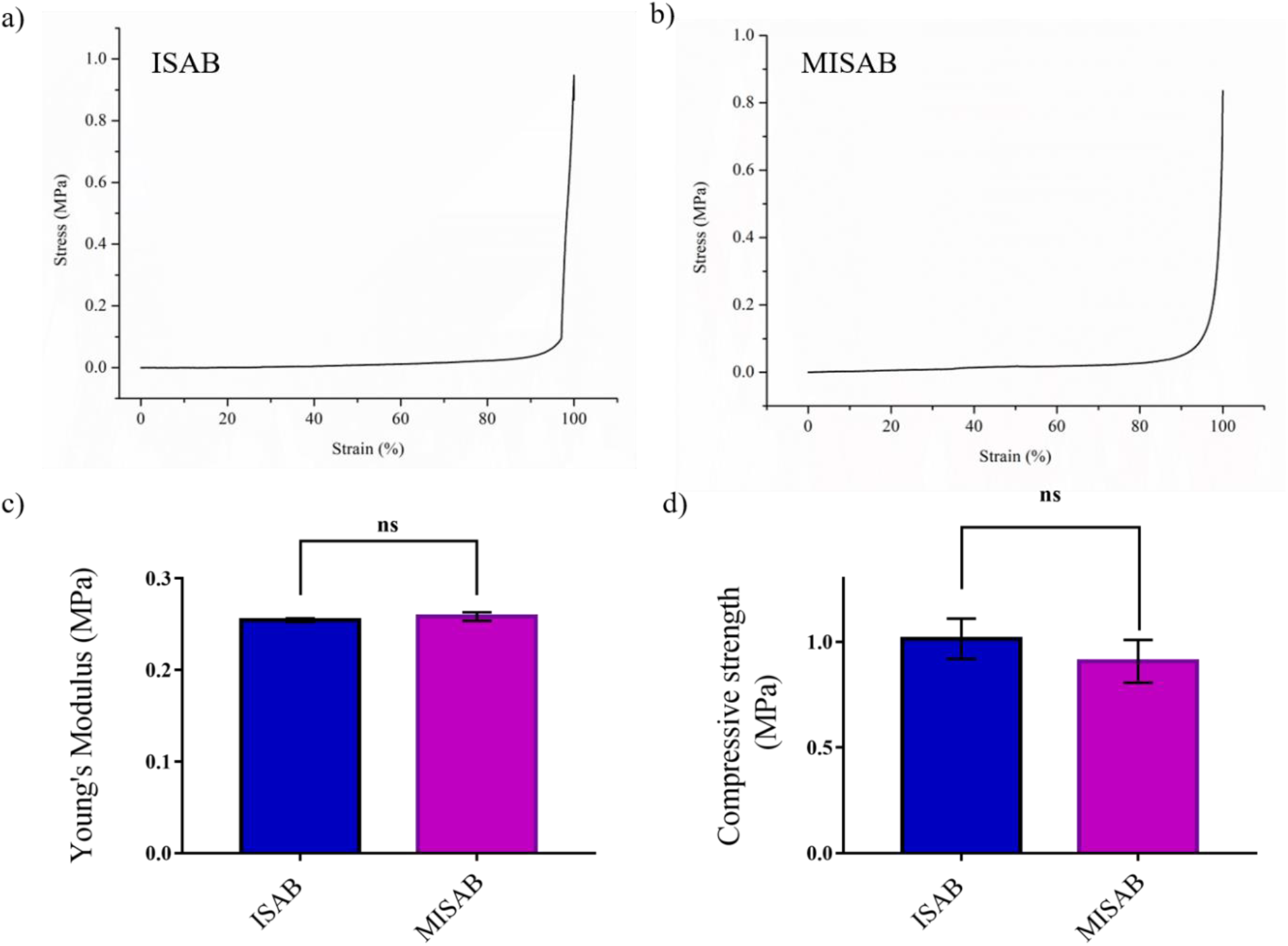
a) & b) Uniaxial compression stress-strain diagram of ISAB and MISAB, respectively b) Young’s modulus of scaffolds d) Compressive strength of scaffolds

**Figure 6:**
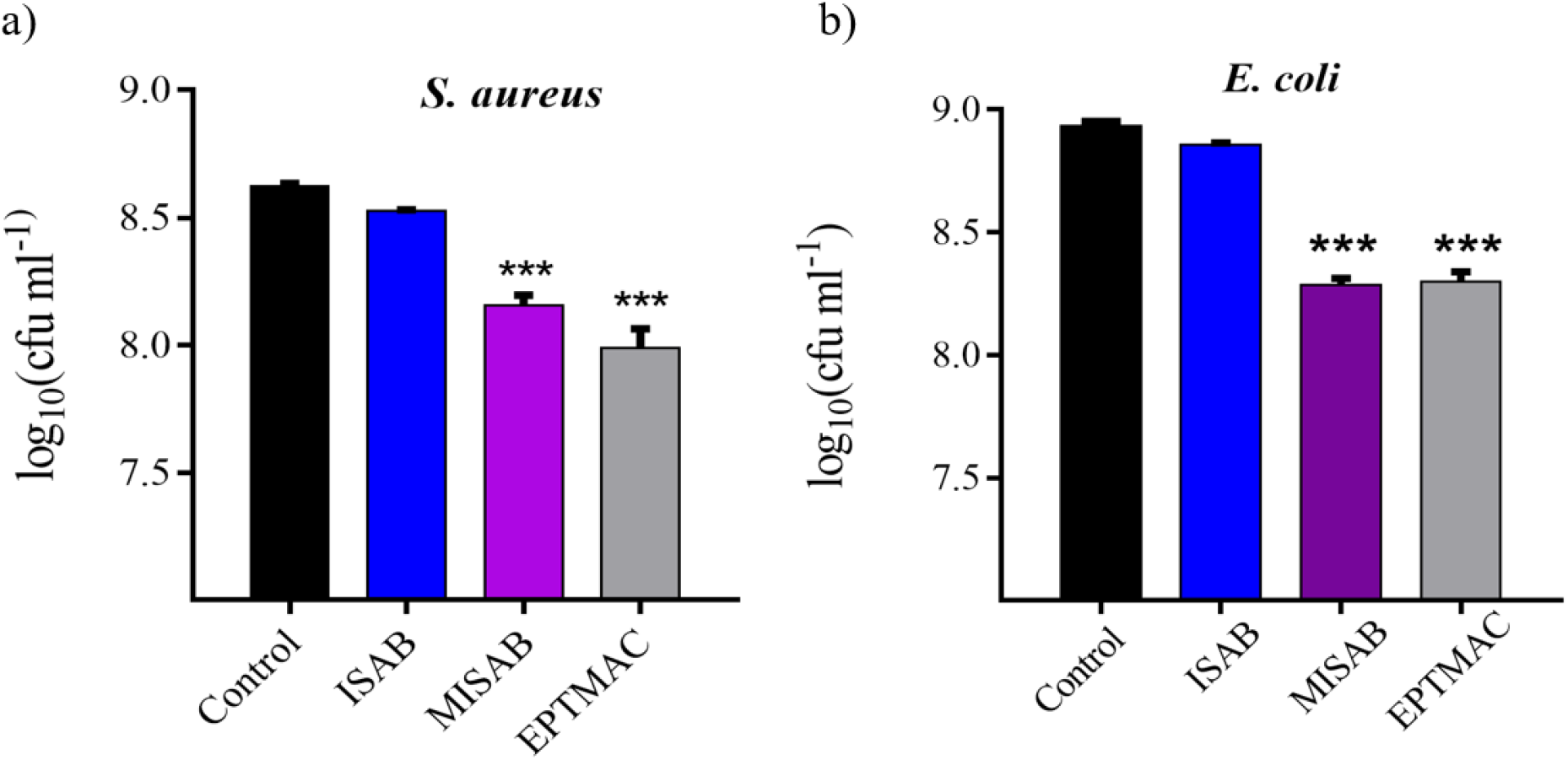
Antibacterial activity of ISAB and MISAB against *S. aureus* and *E. coli* assessed by colony counting assay. Values are the mean±SD (n=3), and the level of significance is denoted as *** (p < 0.001)

### 3.3 Biological Characterization

Antibacterial activity of the ISAB and MISAB scaffolds was evaluated using both gram-negative (*E. coli*) and gram-positive (*S. aureus*) bacteria. The antibacterial activity was measured by counting the number of colonies formed per mL (CFU/mL). MISAB scaffolds showed significant antibacterial activity against *E. coli* and *S. aureus* compared to ISAB scaffolds (p < 0.001) (Figure 6). This activity of MISAB can be attributed to the crosslinking of EPTMAC to Isabgol. Earlier reports have shown that the antibacterial activity of EPTMAC is due to its positive charge, which facilitates its interaction with negatively charged phospholipids on the bacterial membrane and causes membrane disruption [31].

The cell viability of the scaffolds was assessed by the MTT assay (Figure 7a). After 24, 48 and 72 hours of incubation with the scaffold leachate, the MTT assay results indicated that Isabgol crosslinked with EPTMAC showed no significant difference in cell viability compared to the control cells in which no leachate was added. The scaffolds are therefore non-toxic and biocompatible and suitable for use in wound healing applications. Figure 7c shows fluorescence microscope images of control and treated cells stained with SYTO 9/ Propidium iodide. The cells showed normal morphology and most cells stained green indicating that they were alive. Very few cells were dead and stained red with the membrane permeable propidium iodide. The results from the live/dead assay were consistent with the quantitative results obtained in the MTT assay. Scratch wound assay was performed to evaluate *in vitro* wound healing by studying cell migration in the presence of ISAB and MISAB leachate. A scratch was made in a confluent monolayer of fibroblast cells to mimic a wound followed by time-lapse imaging of the cells. Figure 7b shows that the proliferation of cells remained unaffected in the presence of the scaffold leachate. 100% of the wound area was covered after 24 hours in both control and treated wells.

**Figure 7:**
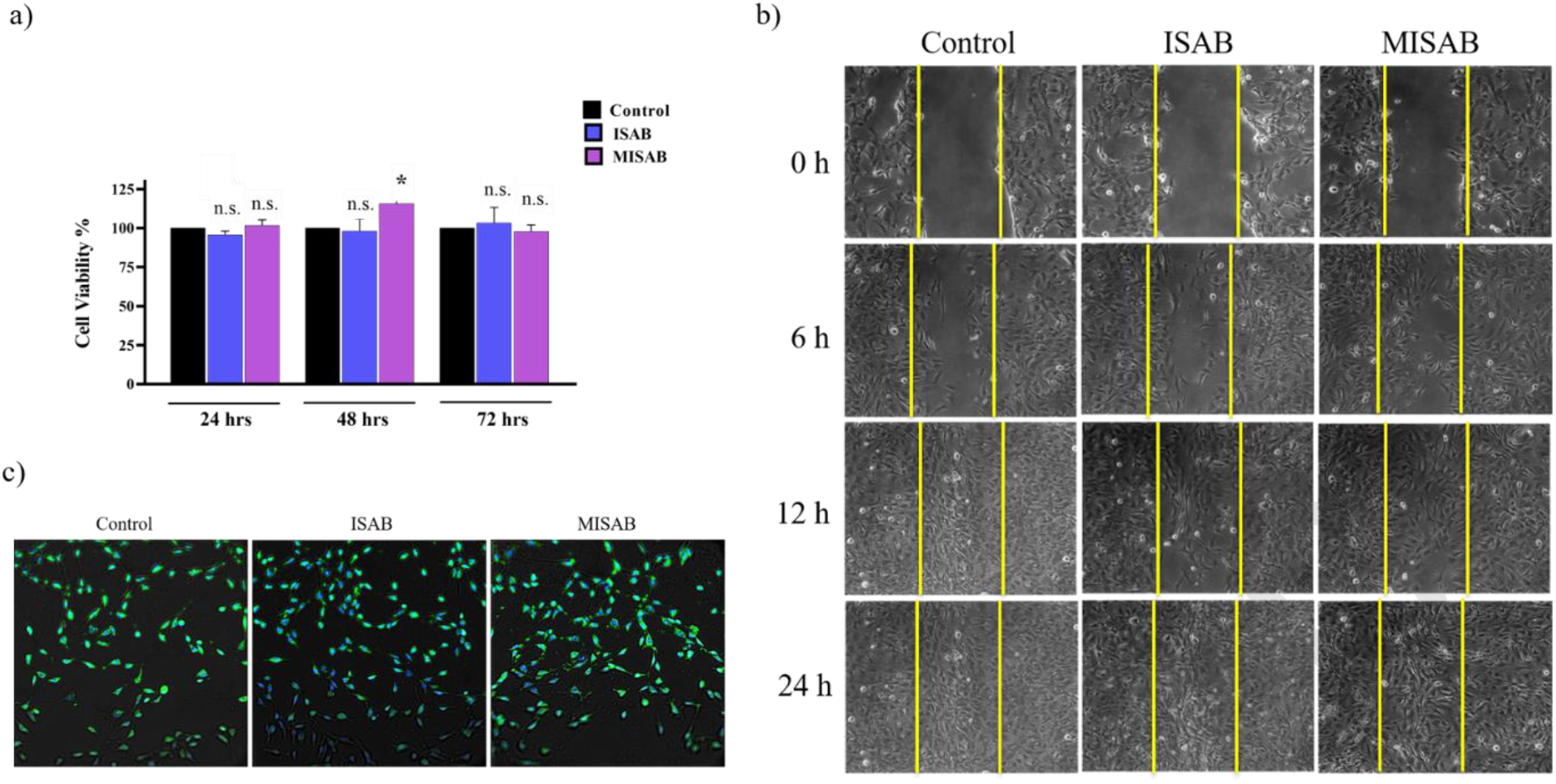
a) : a) Determination of cell viability by MTT assay. Values are expressed as mean values ± SD (*n* = 3); * *p* < 0.05 b) *In vitro* scratch wound healing assay demonstrating the proliferation of cells into the wound area c) Live/dead staining of fibroblast cells treated with ISAB and MISAB scaffold leachate.

An *in vitro* hemolysis assay was performed to assess the hemocompatibility of the scaffolds. The SEM images of the RBCs incubated with the leachate did not indicate any cell lysis (Figure 8a). The morphology of the RBCs in the treated samples was same as the negative control. The positive control with Triton X-100 showed rupture of the RBC membrane and a deviation of shape from the typical biconcave shape of RBCs was observed. Figure 8b shows the release of hemoglobin by the ruptured RBCs in positive control while in the negative control and the treated suspension, the supernatant is clear indicating no rupture of RBCs. Determination of the percentage hemolysis indicated zero lysis in ISAB, 0.24±0.002% lysis in MISAB, and 0.15±0.002% lysis in EPTMAC (Figure 8c).

**Figure 8:**
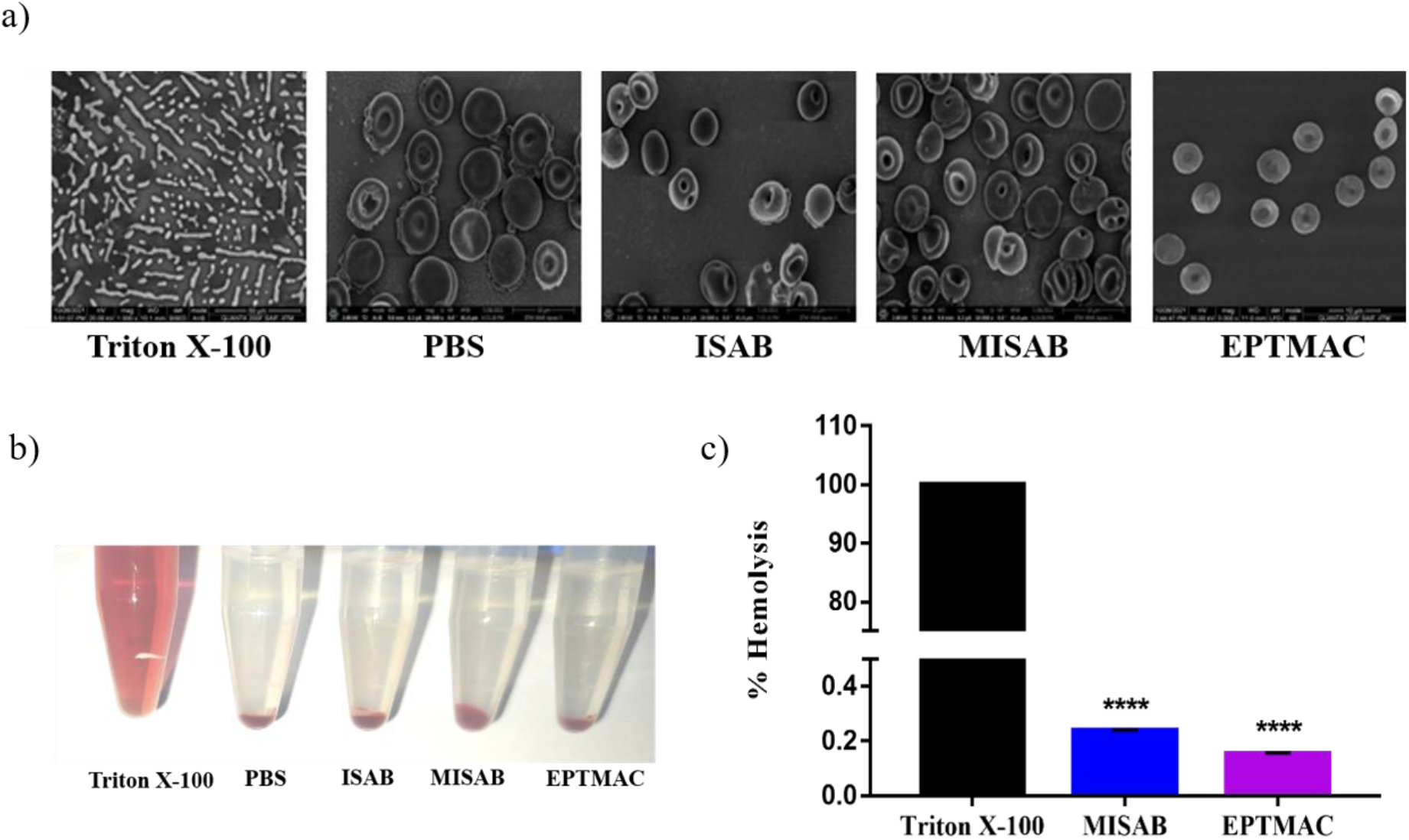
Hemocompatibility study of ISAB and MISAB leachate a) Morphology of RBCs after 4 h incubation with leachate b) Photograph of RBC suspension centrifuged after 8-hour incubation with leachate c) Graph depicting the percentage hemolysis in presence of leachate

## 4. Conclusion

In the present study, we have developed an ingenious and inexpensive wound dressing material that is biocompatible and anti-infective to achieve effective wound management. We have functionalized the natural polymer, Isabgol (Psyllium) with epoxy propyl trimethyl ammonium chloride (EPTMAC), a quaternary ammonium compound. Cationization of the functional groups of the polymer renders antibacterial activity to the material, which can ward-off infections at the wound site. The crosslinking of EPTMAC to Isabgol was performed by modifying the hydroxyl groups on Isabgol via an epoxy ring-opening reaction. The cationized scaffolds were synthesized by lyophilization of the reacted mixture after dialysis. The crosslinking was confirmed by ^13^C NMR spectroscopy and FTIR spectroscopy. A degree of substitution of 60% was obtained as per CHN element analysis. The material was found to be of porous structure by scanning electron microscopy, which will allow the exchange of gases through the material. The material has excellent swelling property with a swelling ratio of 17.4±0.3, aiding in the absorption of wound exudates and maintenance of a moist environment at the wound site. Thermogravimetric analysis showed that the material is stable up to 300°C. From colony forming unit assay, the scaffolds were found to be antibacterial against *Escherichia coli* (Gram negative) and *Staphylococcus aureus* (Gram positive). Cytotoxicity studies performed on 3T3-L1 mouse fibroblast cell line demonstrated no significant reduction in cell viability in the presence of EPTMAC-modified Isabgol scaffolds. An *in vitro* scratch wound assay showed that cell proliferation in the presence of the material was similar to that in the control. The material was proved to be suitable for blood contact application such as wound management through hemocompatibility assays. Therefore, the developed material was found to be biocompatible, anti-infective and suitable for wound dressing applications.

## Acknowledgements

The authors acknowledge funding from the Science & Engineering Research Board (SERB), Department of Science and Technology (DST), Government of India (Project ECR/2017/003064). We thank the Sophisticated Analytical Instrument Facility (SAIF), IIT Madras for SEM facility. Acknowledgments are due to Prof. Nitish Mahapatra for offering their mammalian cell culture expertise and facility.

## Notes

### Competing Interest Statement

The authors have declared no competing interest.

## References

[1] V. Falanga, Wound healing and its impairment in the diabetic foot, Lancet (London, England) 366(9498) (2005) 1736–43.

[2] L. Uccioli, V. Izzo, M. Meloni, E. Vainieri, V. Ruotolo, L. Giurato, Non-healing foot ulcers in diabetic patients: general and local interfering conditions and management options with advanced wound dressings, Journal of wound care 24(4 Suppl) (2015) 35–42.

[3] Z. Xu, S. Han, Z. Gu, J. Wu, Advances and Impact of Antioxidant Hydrogel in Chronic Wound Healing, 9(5) (2020) 1901502.

[4] I.V. Yannas, J.F. Burke, Design of an artificial skin. I. Basic design principles, Journal of biomedical materials research 14(1) (1980) 65–81.

[5] V. Dill, M. Mörgelin, Biological dermal templates with native collagen scaffolds provide guiding ridges for invading cells and may promote structured dermal wound healing, International wound journal 17(3) (2020) 618–630.

[6] X. Feng, X. Zhang, S. Li, Y. Zheng, X. Shi, F. Li, S. Guo, J. Yang, Preparation of aminated fish scale collagen and oxidized sodium alginate hybrid hydrogel for enhanced full-thickness wound healing, International journal of biological macromolecules 164 (2020) 626–637.

[7] X. Du, L. Wu, H. Yan, Z. Jiang, S. Li, W. Li, Y. Bai, H. Wang, Z. Cheng, D. Kong, L. Wang, M. Zhu, Microchannelled alkylated chitosan sponge to treat noncompressible hemorrhages and facilitate wound healing, Nature Communications 12(1) (2021) 4733.

[8] K. Kalantari, E. Mostafavi, B. Saleh, P. Soltantabar, T. Webster, Chitosan/PVA Hydrogels Incorporated with Green Synthesized Cerium Oxide Nanoparticles for Wound Healing Applications, European Polymer Journal 134 (2020) 109853.

[9] S. Shafei, M. Khanmohammadi, R. Heidari, H. Ghanbari, V. Taghdiri Nooshabadi, S. Farzamfar, M. Akbariqomi, N.S. Sanikhani, M. Absalan, G. Tavoosidana, Exosome loaded alginate hydrogel promotes tissue regeneration in full-thickness skin wounds: An in vivo study, Journal of biomedical materials research. Part A 108(3) (2020) 545–556.

[10] Y. Hao, W. Zhao, L. Zhang, X. Zeng, Z. Sun, D. Zhang, P. Shen, Z. Li, Y. Han, P. Li, Q. Zhou, Bio-multifunctional alginate/chitosan/fucoidan sponges with enhanced angiogenesis and hair follicle regeneration for promoting full-thickness wound healing, Materials & Design 193 (2020) 108863.

[11] Z. Hadisi, M. Farokhi, H.R. Bakhsheshi-Rad, M. Jahanshahi, S. Hasanpour, E. Pagan, A. Dolatshahi-Pirouz, Y.S. Zhang, S.C. Kundu, M. Akbari, Hyaluronic Acid (HA)-Based Silk Fibroin/Zinc Oxide Core-Shell Electrospun Dressing for Burn Wound Management, Macromolecular bioscience 20(4) (2020) e1900328.

[12] X. He, X. Liu, J. Yang, H. Du, N. Chai, Z. Sha, M. Geng, X. Zhou, C. He, Tannic acid-reinforced methacrylated chitosan/methacrylated silk fibroin hydrogels with multifunctionality for accelerating wound healing, Carbohydr Polym 247 (2020) 116689.

[13] B. Tao, C. Lin, Z. Yuan, Y. He, M. Chen, K. Li, J. Hu, Y. Yang, Z. Xia, K. Cai, Near infrared light-triggered on-demand Cur release from Gel-PDA@Cur composite hydrogel for antibacterial wound healing, Chemical Engineering Journal 403 (2021) 126182.

[14] X. Zhao, Y. Liang, Y. Huang, J. He, Y. Han, B. Guo, Physical Double-Network Hydrogel Adhesives with Rapid Shape Adaptability, Fast Self-Healing, Antioxidant and NIR/pH Stimulus-Responsiveness for Multidrug-Resistant Bacterial Infection and Removable Wound Dressing, 30(17) (2020) 1910748.

[15] H. Zhao, J. Huang, Y. Li, X. Lv, H. Zhou, H. Wang, Y. Xu, C. Wang, J. Wang, Z. Liu, ROS-scavenging hydrogel to promote healing of bacteria infected diabetic wounds, Biomaterials 258 (2020) 120286.

[16] M. Yin, Y. Wang, Y. Zhang, X. Ren, Y. Qiu, T.S. Huang, Novel quaternarized N-halamine chitosan and polyvinyl alcohol nanofibrous membranes as hemostatic materials with excellent antibacterial properties, Carbohydr Polym 232 (2020) 115823.

[17] J.L. Daristotle, L.W. Lau, M. Erdi, J. Hunter, A. Djoum, Jr., P. Srinivasan, X. Wu, M. Basu, O.B. Ayyub, A.D. Sandler, P. Kofinas, Sprayable and biodegradable, intrinsically adhesive wound dressing with antimicrobial properties, Bioeng Transl Med 5(1) (2019) e10149–e10149.

[18] C.H. Lee, K.S. Liu, C.W. Cheng, E.C. Chan, K.C. Hung, M.J. Hsieh, S.H. Chang, X. Fu, J.H. Juang, I.C. Hsieh, M.S. Wen, S.J. Liu, Codelivery of Sustainable Antimicrobial Agents and Platelet-Derived Growth Factor via Biodegradable Nanofibers for Repair of Diabetic Infectious Wounds, ACS infectious diseases 6(10) (2020) 2688–2697.

[19] C.W. Hicks, E. Selvin, Epidemiology of Peripheral Neuropathy and Lower Extremity Disease in Diabetes, Current diabetes reports 19(10) (2019) 86.

[20] J.F. Kennedy, J.S. Sandhu, D.A.T. Southgate, Structural data for the carbohydrate of ispaghula husk ex Plantago ovata forsk, Carbohydrate Research 75 (1979) 265–274.

[21] T. Ponrasu, P.K. Veerasubramanian, R. Kannan, S. Gopika, L. Suguna, V. Muthuvijayan, Morin incorporated polysaccharide–protein (psyllium–keratin) hydrogel scaffolds accelerate diabetic wound healing in Wistar rats, RSC Advances 8(5) (2018) 2305–2314.

[22] P. Thangavel, R. Kannan, B. Ramachandran, G. Moorthy, L. Suguna, V. Muthuvijayan, Development of reduced graphene oxide (rGO)-isabgol nanocomposite dressings for enhanced vascularization and accelerated wound healing in normal and diabetic rats, J Colloid Interface Sci 517 (2018) 251–264.

[23] I. Shahid Ul, B.S. Butola, Recent advances in chitosan polysaccharide and its derivatives in antimicrobial modification of textile materials, International journal of biological macromolecules 121 (2019) 905–912.

[24] J. Junthip, N. Tabary, M. Maton, S. Ouerghemmi, J.N. Staelens, F. Cazaux, C. Neut, N. Blanchemain, B. Martel, Release-killing properties of a textile modified by a layer-by-layer coating based on two oppositely charged cyclodextrin polyelectrolytes, International journal of pharmaceutics 587 (2020) 119730.

[25] J. Ru, Z. Wang, C. Tong, H. Liu, G. Wang, Z. Peng, Nonleachable Antibacterial Nanocellulose with Excellent Cytocompatible and UV-Shielding Properties Achieved by Counterion Exchange with Nature-Based Phenolic Acids, ACS Sustainable Chemistry & Engineering 9(47) (2021) 15755–15767.

[26] J.A. Sirviö, M.Y. Ismail, K. Zhang, M.V. Tejesvi, A. ÄmmäCläC, Transparent lignin-containing wood nanofiber films with UV-blocking, oxygen barrier, and anti-microbial properties, Journal of Materials Chemistry A 8(16) (2020) 7935–7946.

[27] F. Xie, X. Bian, Y. Lu, T. Xia, D. Xu, Y. Wang, J. Cai, Versatile antibacterial surface with amphiphilic quaternized chitin-based derivatives for catheter associated infection prevention, Carbohydr Polym 275 (2022) 118683.

[28] W. Qiu, Q. Wang, M. Li, N. Li, X. Wang, J. Yu, F. Li, D. Wu, Peptidoglycan-inspired peptide-modified injectable hydrogels with enhanced elimination capability of bacterial biofilm for chronic wound healing, Composites Part B: Engineering 227 (2021) 109402.

[29] A. Mesdaghinia, Z. Pourpak, K. Naddafi, R.N. Nodehi, Z. Alizadeh, S. Rezaei, A. Mohammadi, M. Faraji, An in vitro method to evaluate hemolysis of human red blood cells (RBCs) treated by airborne particulate matter (PM10), MethodsX 6 (2019) 156–161.

[30] Y. Liang, X. Zhao, T. Hu, Y. Han, B. Guo, Mussel-inspired, antibacterial, conductive, antioxidant, injectable composite hydrogel wound dressing to promote the regeneration of infected skin, Journal of Colloid and Interface Science 556 (2019) 514–528.

[31] Y. Jiao, L.-n. Niu, S. Ma, J. Li, F.R. Tay, J.-h. Chen, Quaternary ammonium-based biomedical materials: State-of-the-art, toxicological aspects and antimicrobial resistance, Progress in Polymer Science 71 (2017) 53–90.

## Web reference

36. National Center for Biotechnology Information (2022). PubChem Compound Summary for CID 18205, 2,3-Epoxypropyltrimethylammonium chloride. Retrieved February 7, 2022 from https://pubchem.ncbi.nlm.nih.gov/compound/2_3-Epoxypropyltrimethylammonium-chloride.

